# Allometric studies on human- and pig-sized Delphinidae to understand resting metabolic rates with respiratory measurements

**DOI:** 10.1101/2025.01.22.633296

**Authors:** Ippei Suzuki, Yasuaki Niizuma, Daiki Inamori, Yurie Watanabe, Katsufumi Sato, Kagari Aoki

## Abstract

Exponential relationships between basal metabolic rate (BMR) and body mass are known in both terrestrial and marine mammals, with elevated BMR in the latter. However, it remains unclear whether the exponential relationship in marine mammals is different from that in terrestrial mammals. Metabolic rates of animals while resting at the water surface (resting metabolic rates: RMRs) were measured by breath-by-breath respirometry in two species of Delphinidae from temperate regions to elucidate the relationship between RMR and body mass in pantropical spotted dolphins (*Stenella attenuate*) and Risso’s dolphins (*Grampus griseus*) with body masses ranging from 71 to 386 kg. The RMRs of rough-toothed dolphins (83–123 kg), common bottlenose dolphins (143–220 kg), and killer whales (3,364–5,318 kg) were obtained from the literature for allometric analyses. Measured RMRs were 344 ± 39 and 1398 ± 160 ml O_2_ min^-1^, respectively, which were 1.2–2.1 times higher than those predicted for similar-sized terrestrial mammals. A linear mixed-effects model showed the allometric mass exponent of 0.65 (95% CI: 0.54–0.76), which was closer to the 2/3-power of body mass than that of the widely observed 3/4-power commonly reported across a broad range of taxa. The new scaling relationship observed in the Delphinidae indicates that larger dolphins require less energy in aquatic environments, which is consistent with the theory of heat loss associated with a narrower surface area relative to larger volumes. This study suggested a different scaling relationship between aquatic and terrestrial mammals.

## INTRODUCTION

The cost of living is essential information describing the baseline energetic demands for the survival of organisms and is a useful metric in comparative physiology to understand the difference in organisms according to their surrounding environments. The energy expenditure required to maintain homeostasis without any active motion is defined as the basal metabolic rate (BMR) in terrestrial mammals and the resting metabolic rate (RMR) in marine mammals, to clarify that the metabolic rate is measured while resting on the water surface (Williams, 2022). Historical allometric studies have found a scaling relationship between BMR and body mass (*M*_b_), which can be expressed as BMR = *aM*_b_*^c^*, where *c* is the allometric mass exponent, and *a* is a constant (Rubner, 1883). The range of values of allometric mass exponents has been debated for over a century, beginning with Rubner’s surface hypothesis suggesting a 2/3-power (Rubner, 1883) and followed by Kleiber’s law proposing a range of 0.734–0.75, which is famous for scaling from mice to elephants (Kleiber, 1932; Brody, 1945: Kleiber, 1961). A study also suggested that defining the universal allometric mass-exponent would not be realistic because each animal adapts its morphology and physiology to each realm, while physical rules are dominated by body size, which is important for the ‘default,’ or standard, to conduct inter-species comparative biology (Lindstedt and Hoppeler, 2023).

Quantification of heat production is theoretically the most direct measurement of metabolism (direct calorimetry), although it is essentially impossible to apply to marine mammals because these animals exclusively live in aquatic environments. Collecting breaths to measure the rate of oxygen consumption (indirect calorimetry) was alternatively used for measuring RMR in marine mammals (Kooyman et al., 1973; Matsuura and Whittow, 1973), which discovered that RMR in marine mammals are 0.2–3.6 multiples of Kleiber’s equation (reviewed in Noren and Rosen, 2023). Higher RMR in marine mammals was initially attributed to high metabolic demands for thermoregulation during exposure to cold climates in high latitudes or water environments with 25 times higher thermal conductivity than in air (Ridgway, 1972). This explanation applies to newborn or juvenile individuals with insufficient adipose tissue (Whittow, 1987), but not to mature animals, which possess thick layer of blubber for insulation, although the full mechanism of behavioral and physiological thermoregulation remains to be investigated (Favilla et al., 2022; Sakai et al., 2024). The surface-to-volume ratio (SVR) is a crucial factor in aquatic environments because heat loss is proportional to the surface area (Davis, 2019). Larger animals had a lower SVR than that of smaller animals, leading to reduced heat loss. This tendency for body size to increase in colder regions among closely-related species is known as the Bergmann’s rule (Mayr, 1963).

BMR measurements require several criteria: a non-growing adult individual in a resting condition, which is postabsorptive in a thermoneutral environment (Kleiber, 1961). In fact, metabolic rates were affected by digestion process and growth stages. For example, 63% of increase in metabolism from the postabsorptive to postprandial condition was reported in juvenile South American fur seals (Dasis et al., 2014) and 29-53% increase of metabolic rates after ingestion were observed in adult bottlenose and rough-toothed dolphins (van der Hoop et al., 2014; Fahlman et al., 2024). Early metabolic measurements on marine mammals did not always meet these criterions because it had been challenging to unify experimental conditions due to difficulties to control animals’ behavior without feeding and physical restrictions to fit large and matured animals to experimental facilities. These historical challenges have been also considered as partial reasons of elevated RMR in marine mammals compared with similar-sized terrestrial mammals (Noren and Rosen, 2023). Recent studies which met the Kleiber’s criteria also reported that measured RMRs were not as high as previously reported in common bottlenose dolphins (Allen et al., 2022) and for warm-water species in Hawaiian monk seals (John et al., 2021). Combining these historic challenges and recent findings, measuring the RMRs of warm-water species with a wide range of body masses would provide a better understanding of the metabolic rates of marine mammals compared to those of terrestrial mammals.

The Delphinidae is one of the most speciose and successful families among marine mammals with more than thirty recognized extant species and wide habitat distributions from tropical to polar regions (LeDuc, 2009). Their body size ranges from 40 kg to >5,000 kg, which wide range of body size is ideal for conducting allometric study, but knowledge on their metabolisms is limited for only a few species due to the difficulties of conducting metabolic measurements under the Kleiber’s criteria. In this study, we conducted respirometry measurements under the Kleiber’s criteria on two species of Delphinidae, the pantropical spotted dolphin (*Stenella attenuata*) and Risso’s dolphin (*Grampus griseus*), with body masses ranging from 71 kg to 386 kg. Despite their broad distribution across the temperate zones of the world’s oceans, surprisingly little is known about the metabolic traits of these species. In addition, advances in sensory technologies have enabled us to directly measure instantaneous changes in oxygen and tidal volumes from dolphin blows, without collecting a large volume of each blow (Fahlman et al., 2015; hereafter called “breath-by-breath respirometry”), which mitigates the difficulties in conducting respiratory experiments on medium- and large-sized dolphins. Historically, metabolic rates in captive dolphins have been measured with a floating dome on the water surface where dolphins were trained to breath inside the dome, which is called open-flow respirometry (Fedak et al., 1981; Williams et al., 1993). A side-by-side comparison between open-flow and breath-by-breath respirometry on unfasted metabolic rates confirmed similar data quality between the two methods using the same captive bottlenose dolphins with similar body sizes in Hawaii (van der Hoop et al., 2014; Fahlman et al., 2015; Supplementary Figure S1). We applied the breath-by-breath respirometry to our study animals.

The aim of this study is to fill gaps in knowledge about metabolic traits of small- and large-sized dolphins in Delphinidae for better understanding on the relationship between RMRs and body mass in aquatic environments. Considering the effect of heat loss in regard to SVR, larger dolphins are expected to have lower mass-specific RMRs than those of smaller dolphins. Using captive animals with daily training in the aquarium enabled the design of consistent experimental conditions in terms of growth stages and digestion processes by controlling the timing of feeding.

## MATERIALS AND METHODS

### Study individuals

This study aimed to elucidate the resting metabolic rate and respiration of delphinids which are under human care. All procedures, animal husbandry, and management were performed under the careful supervision of veterinarians and veterinary nurses at the Whale Museum and Aquarium (Wakayama, Japan). Respirometry data were collected from adult female Risso’s dolphins (animal identification [ID]: gg_cf and gg_nf) in April-October 2021 and from one adult female and one adult male pantropical spotted dolphin (ID st_ml and ID st_mr; Table 1) in August-September 2022. The females were neither pregnant nor lactating during the study period.

**TABLE 1.** Biological information and resting metabolic rate (RMR) of two adult pantropical spotted dolphins and an adult Risso’s dolphin. The body length and girth at the anterior insertion of dorsal fin were measured in August 2022 for the pantropical spotted dolphins, and in July and October 2021, for the Risso’s dolphin. Species name, animal identification (ID), sex, body mass (*M*_b_), body length, girth length, estimated age (Age), display periods, number of trials, total number of breaths, RMR, and Kleiber ratio. Mean ± SD.

| Species | ID | Sex | $M_b$<br>(kg) | Body<br>length<br>(cm) | Girth<br>length<br>(cm) | Age**<br>(year) | Display<br>periods**<br>(year) | No. of<br>trials | Total no. of<br>breaths | RMR<br>(mL O<sub>2</sub> min<sup>-1</sup>) | Kleiber ratio |
| --- | --- | --- | --- | --- | --- | --- | --- | --- | --- | --- | --- |
| Pantropical<br>spotted<br>dolphin | st_mr | Female | 72 | 200 | 94 | 20 | | 4 | 68 | 334 ± 37 | 1.3 ± 0.1 |
|  | st_ml | Male | 71 | 193 | 96 | 14 | 10 | 1 | - | 384 | 1.6 |
| Risso's dolphin | gg_cf | Female | 315 ± 9* | 305 ± 4 | 151 ± 2 | 16 | 11 | 3 | 101 | 1,418 ± 189 | 1.9 ± 0.2 |
|  | gg_nf | Female | 386 | 291 | 178 | 17 | 13 | 1 | 30 | 1,335 | 1.5 |

\*Body mass was estimated from the body length and girth based on Tobayama (1999).
\*\*Estimated age and display periods during measurement period (2022 for pantropical spotted dolphins and 2021 for a Risso's dolphin) were shown.

Pantropical spotted dolphins were housed in an outdoor swimming pool with a shelter (length: 7 m; width: 15 m; depth: 3–4 m). The pool was connected to a nearby bay to ensure constant exchange of seawater between the pool and bay. Risso’s dolphins were housed in a large natural semi-enclosed area with natural furniture and shelter (50 × 210 × 2–6 m, length × width × depth). Net pens (length: 12 m; width: 12 m; depth: 3–4 m) were temporarily employed for healthcare and experimental procedures. The enclosure was surrounded by hills and forests, which provided a shaded environment.

Individuals were provided with a mixed diet consisting of mackerel and squid, supplemented with vitamins multiple times a day. Table 1 summarizes the ID, body length, body mass, sex, and estimated ages of the four study individuals.

### Respiratory data sampling

We measured the respiratory flow of the individuals. The animals were fasted over-night to make their digestion process in postprandial to meet the Klieber’s criteria for measuring RMRs (Pederson et al., 2020; Allen et al., 2022). A previous study reported that excretion of red-stained prey in captive Indo-Pacific bottlenose dolphins (*Tursiops aduncus*) first observed in their feces after the latest of six hours (Takahashi et al., 2021). We believe that >12 hours of over-night fasting in this study also satisfied the criteria to make the animals postprandial.

Respiratory measurements were conducted early in the morning before the aquarium’s opening hours to minimize the influence of visitors, which timing also coincides with a typical resting period for wild dolphins (Vermeulen et al., 2015). Animals were kept at stationary positions using slings for the health measurement and experiments. The slings were tailored to fit the size of each individual and used for health measurements. The experiment was not conducted if the animal was hesitant to rest on the sling. Consequently, ID st_ml was measured only once. In all the other cases, the individuals were calm during the experiment.

The respiratory measurements were conducted based on the methods described by Fahlman et al. (2015) and Allen et al. (2022), briefly summarized below. The respiratory flow was measured using a custom-made Fleisch-type pneumotachometer (Adm + Engineering, Valencia, Spain, Fig. 1A) equipped with a low-resistance laminar flow matrix (Z9A887–2, Merriam Process Technologies, Cleveland, OH, USA). A differential pressure transducer (Spirometer Pod, ML 311, ADInstruments, Colorado Springs, CO, USA) was connected to the pneumotachometer with two firm-walled and flexible tubes, each with a length of 340 cm and 1.5 mm and 2.0 mm inner diameter for spotted and Risso’s dolphins, respectively (Fig. 1B). The differential pressure transducer was connected to a data acquisition system (PowerLab 4/26, ADInstruments), and the data were captured at 400 Hz and displayed on a laptop computer running LabChart (v. 8.1, ADInstruments). Differential pressure was used to determine the flow rate and was calibrated using a 7.0 liter calibration syringe (Series 4900, Hans-Rudolph Inc., Shawnee, KS, USA). The calibration factor for flow rate was calculated separately for inspired and expired flows, which was relatively consistent across trials for each individual. Normal breathing was defined as respiration that began with exhalation, followed by immediate inhalation (Fahlman et al., 2020; Fig. 2).

**FIGURE 1.**
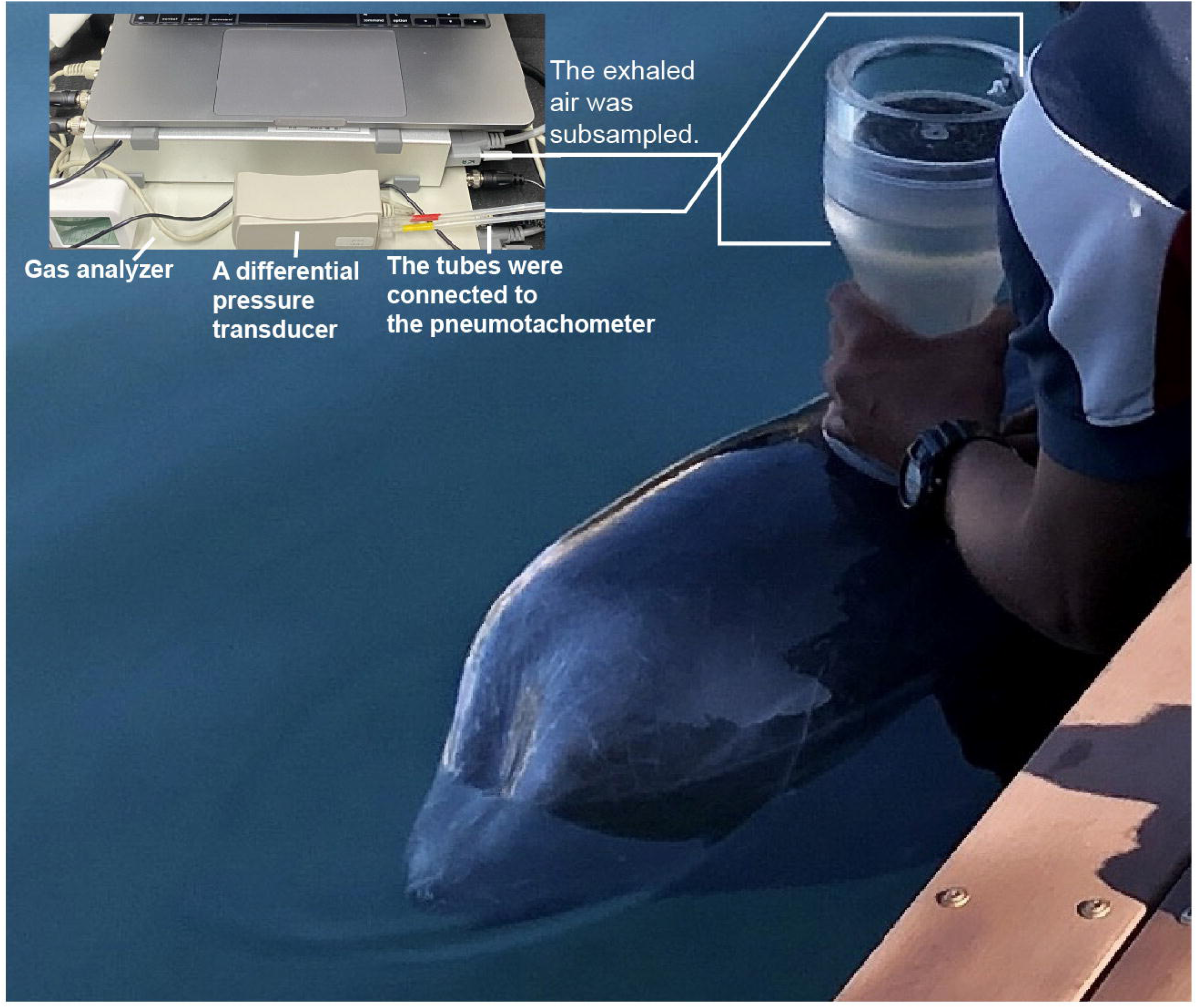
Respiration measurement of a Risso’s dolphin (ID: gg_cf). (**A**) A custom-made Fleisch-type pneumotachometer (adm+ engineering, Valencia, Spain) was attached to the dolphin’s head during the experiment. (**B**) A differential pressure transducer (Spirometer Pod, ML 311, ADInstruments, Colorado Springs, CO, USA) was connected to the pneumotachometer to measure flow rate. The below samples were sub-sampled via a port for gas concentration analysis (O_2_ and CO_2_).

**FIGURE 2.**
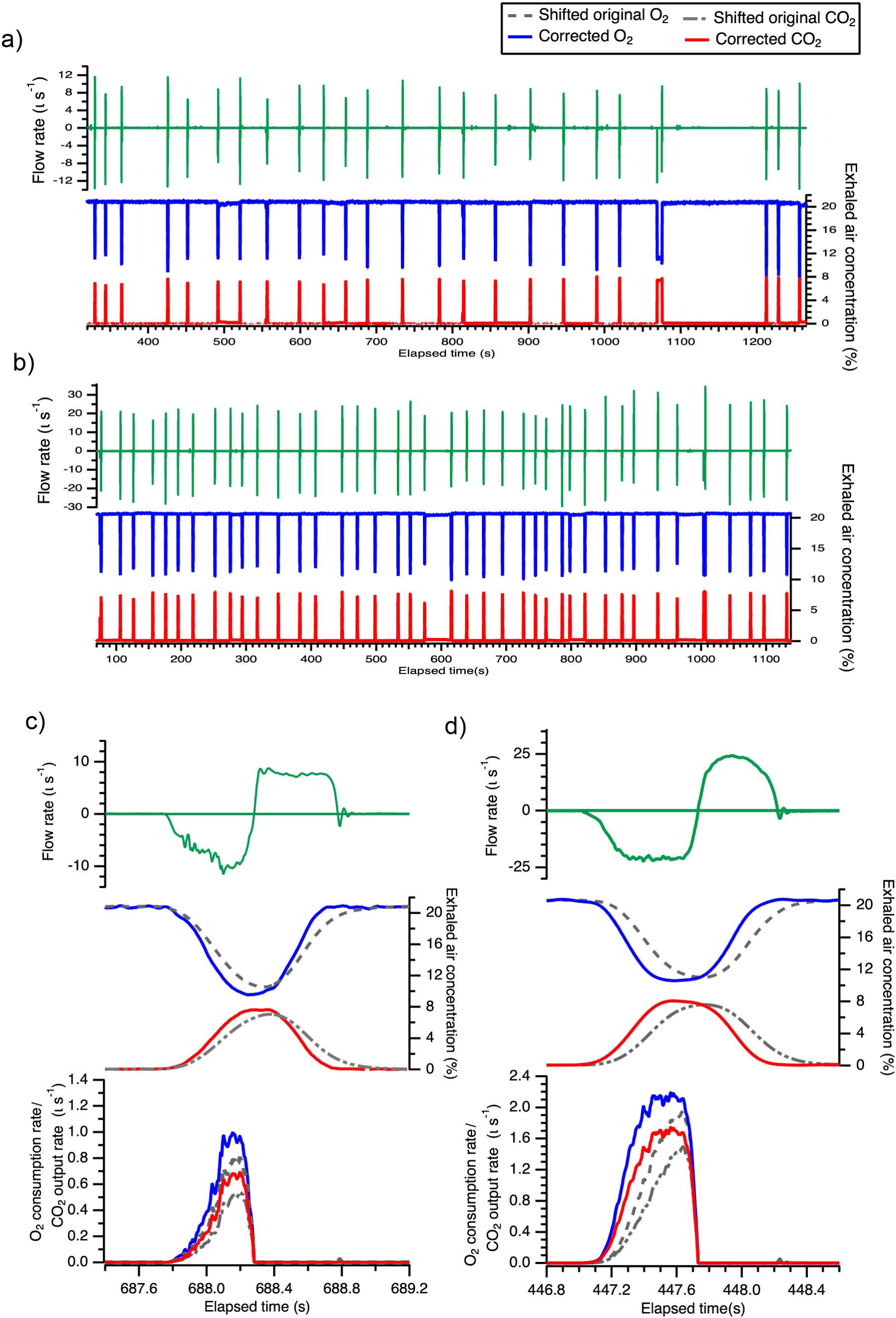
Examples of respiratory time series data. Negative flow rate values represent exhalation, while positive values represent inhalation. **A, B**) Pantropical spotted dolphin: (**A**) an entire trial and (**B**) an enlarged view of a single breath. **C, D**) Risso’s dolphin: (**C**) an entire trial and (**D**) an enlarged view of a single breath. To account for waveform distortion and signal delay caused by the tubing and gas analyzer response time, O_2_ concentration was adjusted using the method described by Allen et al. (2022).

The exhaled air was subsampled via a port in the pneumotachometer and passed through the same type of tubing used for measuring respiratory flow at a flow rate of 200 ml min^-1^. This was followed by a firm-walled, flexible tubing and a 30 cm length of 1.5 mm inner diameter Nafion tubing, to a fast-response gas analyzer (ML206, ADInstruments, with a 95% response time below 200 ms at a flow rate of 200 ml min^-1^). The gas analyzer was connected to a data acquisition system and sampled at 400 Hz. Two-points calibrations for both O_2_ and CO_2_ concentrations were performed daily before the first trial using ambient air (20.9% O_2_, 0.04 % CO_2_) and commercially available calibration gas (a mixture of 5.1% O_2_, 5.2% CO_2_, and 89.7% N_2_; Saisan Co., Ltd., Saitama, Japan). Ambient air was also used to verify the calibration after each trial. Considering cetaceans’ high exhalation rate and to compensate as much as possible for the oxygen transducer’s response time, the rigid Teflon tube was designed with the smallest possible inner diameter (1.5-2.0 mm) and the shortest possible length (340 cm) to measure O_2_ and CO_2_ concentrations. The water temperatures during the trials were 28.4 ± 0.9°C, 26.7 ± 1.5°C, 17.6 °C for two pantropical spotted dolphins (*n* = 5 trials), the Risso’s dolphin (ID gg_cf, *n* = 3 trials) and ID gg_nf (*n* = 1 trial), respectively. The dolphins were desensitized to the pneumotachometer for at least 1 month before the study.

### Resting metabolic rate

The resting metabolic rate (ml O₂ min⁻¹) for each trial was calculated based on Allen et al. 2022. To correct for the delay in subsampled exhaled air reaching the gas analyzer, gas signals were time-shifted to align with the flow rate. All gas volumes were converted to standard temperature and pressure, dry (STPD).

As explained by Allen et al. (2022), the tube and gas analyzer response times cause a distortion in the shape and a delay in the measured signal. The distortion of both gas concentration signals was corrected using a two-exponent equation based on Allen et al. (2022):

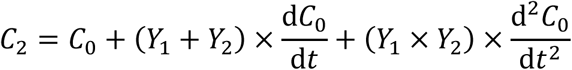

where, *C*_2_ is the corrected signal, *C*_0_ is the low-pass filtered (5 Hz cut-off) original signal, *Y*_1_ and *Y*_2_ are constants, and d*C*_0_/d*t* and d^2^*C*_0_/d*t*^2^ are the first and second derivatives of *C*_0_, respectively. We used *Y*₁ = 0.07 and *Y*₂ = 0.08 to obtain appropriate *C*₂ values for both O₂ and CO₂ in this subset of exhales and inhales. The oxygen consumption rate and CO₂ output rate were calculated by multiplying the corrected O₂ and CO₂ values by the expiratory flow rate, respectively, and the totals were summed over the entire trial.

### Metabolic rates from literatures

In addition to the RMRs measured for the two species in our experiments, previously reported RMRs from common bottlenose dolphins (*Tursiops truncatus*) and rough-toothed dolphins (*Steno bredanensis*) were used for allometric analysis (Table S1). These studies measured RMRs using identical respiratory systems, and our experimental conditions were designed to be consistent with these studies in terms of animal growth stages and digestion processes. All individuals in the literature were mature and the females were not pregnant. One RMR reported in common bottlenose dolphins (9FL3 in Table 1 in Allen et al., 2022) was excluded from our analysis because this particular individual was wild. Furthermore, RMRs of adult killer whales (*Orcinus orca*) in postabsorptive condition were also cited from literatures, which were measured by either a meteorological balloon or open-flow respirometry (Kriete, 1994; Worthy et al., 2013). RMRs measured by open-flow respirometry in common bottlenose dolphins were also cited from a previous study (Yeates and Houser 2008, John et al., 2024; Table S1). The ratio to the Kleiber’s curve (Kleiber ratio) was calculated by dividing measured RMRs by the predicted BMR in terrestrial mammals which is obtained by the following equation in kcal day^-1^: BMR = 70.0 *M* ^0.75^ (Kleiber, 1961).

### Statistical analysis

A linear mixed effects model (LMM) was used with nested random effects of species names to account for differences in individual species and predict the RMR in Delphinidae. Both RMRs and body mass were log10-transformed (Glazier, 2021). The models were fitted to the R statistical computing software (R Core Team, 2021; RStudio Team, 2021) using the *lme4* package (Bates et al., 2015). Mean ± SD are shown.

## RESULTS

The average RMR in the pantropical spotted dolphins was 344 ± 39 ml O_2_ min^-1^ (measurement duration, 877 ± 322 s, *n* = 5 trails, see the Supplemental table for details of each trial) and that in the Risso’s dolphin was 1,398 ± 160 ml O_2_ min^-1^ (measurement duration, 978 ± 223 s, *n* = 4 trails, Table 1). The model in the LMM is the following equation with the fixed effect of log10-transformed body mass:

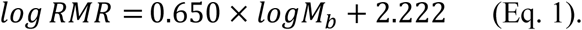

A range of the 95% confidence interval of allometric mass exponent was 0.538-0.762 (Fig. 3). The ratio of measured RMR to that of the Kleiber’s terrestrial mammals were 1.5–2.2 (Table 1).

**FIGURE 3.**
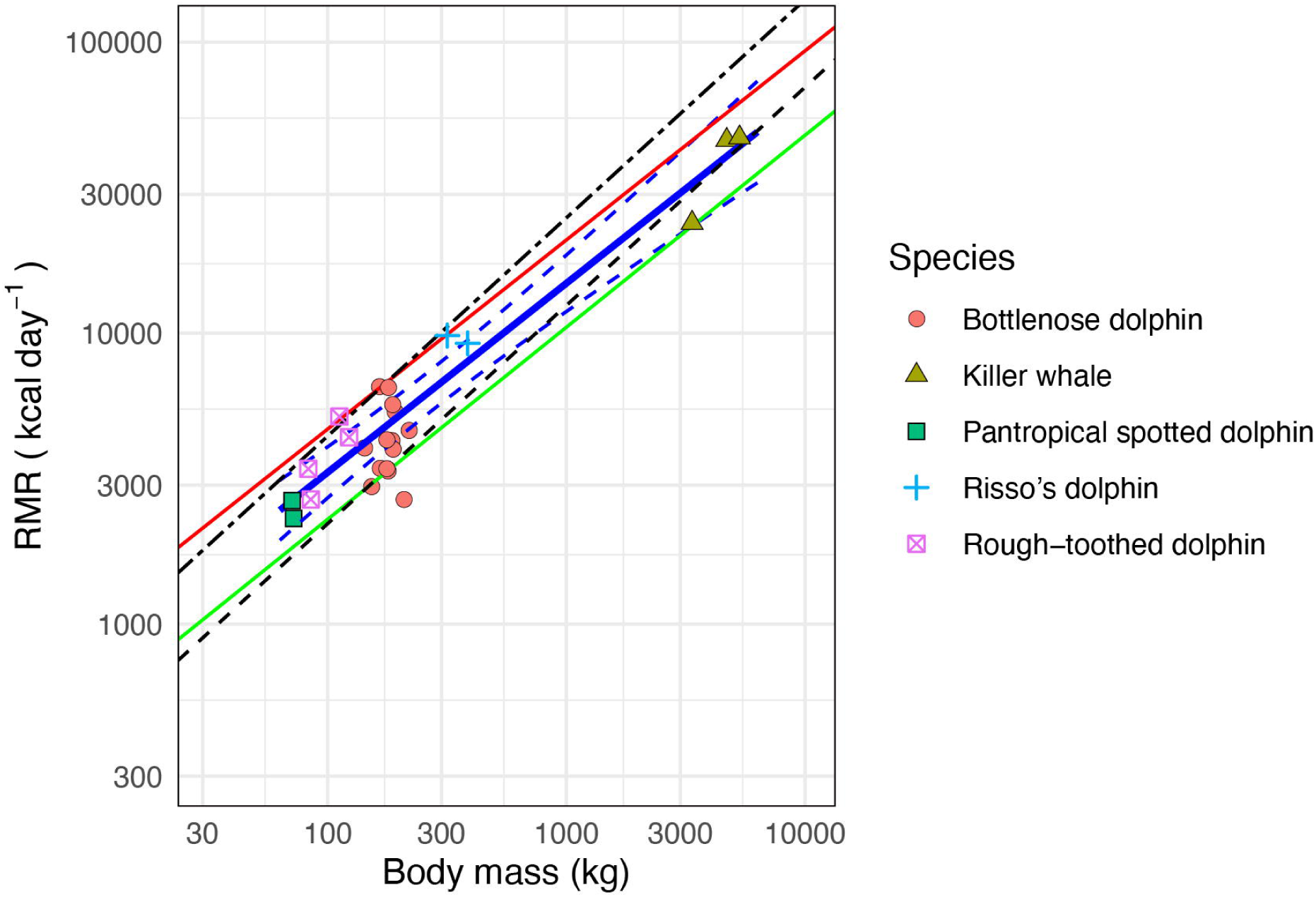
Allometric scaling of resting metabolic rate (RMR) in five species of Delphinidae. The blue solid and dashed lines represent the estimated regression line and the 95% confidence intervals from this study. The black dashed line represents Kleiber’s law, the black dot-dashed line represents twice the value of Kleiber’s law. The red line represents the regression from John et al. (2024) based on Williams et al. (2001), and the green line represents the regression from He et al. (2023).

For the pantropical spotted dolphin (ID st_mr), inspiratory tidal volume (*V*_Tinsp_), the maximum inspired flow rate (*V*_insp_), expiratory tidal volume (*V*_Texp_), the maximum expiratory flow rate (*V*_exp_), end-expired O_2_ and CO_2_, and breath frequency at rest were 10.6 ± 2.0 liters s⁻¹, 3.4 ± 1.5 liters, 10.6 ± 2.0 liters s⁻¹, 2.9 ± 1.2 liters, 9.1 ± 1.3%, 7.9 ± 1.3%, and 2.9 ± 2.7 breath min⁻¹, respectively (*n* = 68 breaths, Table S2). For the Risso’s dolphin (ID gg_cf), *V*_insp_, *V*_Tinsp_, *V*_exp_, *V*_Texp_, end-expired O_2_ and CO_2_, and breath frequency at rest were 24.0 ± 3.3 liters s⁻¹, 9.3 ± 1.8 liters, 24.2 ± 5.3 liters s⁻¹, 9.1 ± 2.1 liters, 11.8 ± 1.2%, 7.5 ± 0.6% and 2.9 ± 4.5 breath min⁻¹, respectively (*n* = 101 breaths, Table S2). For the Risso’s dolphin (ID gg_nf), *V*_insp_, *V*_Tinsp_, *V*_exp_, *V*_Texp_, end-expired O_2_, and breath frequency at rest were 27.1 ± 2.3 liters s⁻¹, 12.2 ± 1.8 liters, 21.8 ± 3.8 liters s⁻¹, 9.4 ± 2.6 liters, 12.1 ± 1.0% and 2.3 ± 0.7 breath min⁻¹, respectively (*n* = 21 breaths, Table S2). Only normal breaths were included in above calculation; hesitant breaths and multiple exhalations before inhalation were excluded.

## DISCUSSION

This study revealed a relationship between the RMR and *M*_b_ of Delphinidae, with an allometric mass exponent of 0.650 (Fig. 3). This value was closer to the 2/3-power of body mass than that of the widely observed 3/4-power commonly reported across a broad range of terrestrial taxa. The lower allometric mass-exponent implies that the metabolic demand for homeostasis in larger animals is lower than that in smaller animals in Delphinidae than that in terrestrial mammals, which is consistent with the theory of thermoregulation because larger animals lose heat more slowly than smaller animals in aquatic environments due to their narrower surface area exposed to water relative to its large volume, as proposed in previous studies (Whittow, 1987; Davis 2019; Williams 2022; He et al., 2023). For example, the Kleiber ratio of a 300 kg Delphinidae was 1.35, while that of a 70 kg dolphin was 1.56, based on the results of the LMM (Eq. 1). Surface area is a product of *L*^2^, where *L* is the length, and its exponential relationship with 2/3-power of *M*_b_ is widely accepted in animals (Schmidt-Nielsen, 1984). An earlier study conducted allometric comparisons including a variety of families in marine mammals from Mustelids, the smallest marine mammals (*M*_b_ < 50 kg), to Balaenopteridae, the largest marine mammals (*M*_b_ > 140,000 kg, Williams, 2022). The present study compared the RMRs between two species of Delphinidae with those in mature individuals (range of *M_b_*: 71–386 kg) in temperate waters and obtained a similar allometric tendency, although Kleiber’s curve was within the confidence interval (Fig. 3). Marine mammals adapted their bodies with some of the following strategies against heat loss to live in aquatic environments where the thermal conductivity is high: 1) a thick layer of blubber, 2) a high density of fur, and 3) high heat production in skeletal muscles. The use of a layer of adipose tissue as insulation is one strategy against heat loss, as most marine mammals do; however, having too much blubber would result in increases in both drag and locomotor costs, as reported in a kinematic study of pregnant dolphins (Noren et al., 2011). In addition to having the densest fur in the kingdom Animalia (Kitchener et al., 2017), higher heat production in locomotor muscles via mitochondrial proton leakage has been reported in the smallest marine mammal species, sea otters (Wright et al., 2021), but application of the same mitochondrial mechanism to other marine mammals remains to be investigated (Favilla et al., 2022; Williams, 2022). The availability of diverse body sizes in Delphinidae provided a different relationship between RMR and *M*_b_ than that in terrestrial mammals, which tended to be similar to the relationship between surface area and *M*_b_ This study also reported a Kleiber ratio ranging from 1.2–2.1 in the pantropical spotted and Risso’s dolphins (Table 1). Previous studies showed 1.1-3.5 times higher RMR in odontocetes than the predicted BMR in terrestrial mammals of similar body mass (Noren and Rosen, 2023), while the maximum ratio of measured RMRs in the present study in Delphinidae was less than 2.1 multiples of Kleiber’s BMR. According to the LMM results, smaller dolphins were expected to have higher metabolic rates than that of Kleiber’s BMR, and the ratio in pantropical spotted dolphins (approximately 70 kg) was less than 1.5. The lower ratio of the pantropical spotted dolphins in the current study could be attributed to their adaptation to warm-water environments. This species is distributed from temperate to tropical regions, where the surface water temperature is above 25°C (Perrin, 2009). While the exact thermoneutral zone of this species is not known, our animals were maintained in water (26.7–28.7°C) similar to their natural habitat and we believe that measured RMRs represent basal levels of metabolism for this species. Warm-water marine mammals tended to have lower Kleiber ratio which was 1.2 in Hawaiian monk seal (John et al., 2021) and 0.2–0.7 in manatees (Gallivan and Best, 1980; Irvine, 1983; John et al., 2021). Comparisons of the conductivity and thickness of blubber between tropical and temperate species of similar-sized dolphins showed 200% higher conductivity and 50% less thickness of blubber in tropical dolphins, which supported less physiological demands for thermoregulations in warm-water environments (Worthy and Edwards, 1990). In addition to warm water adaptations, the lowest ratios observed in sirenians were considered to be due to the different strategies in the digestion process between herbivores and carnivores (John et al., 2021). Therefore, slightly higher, but not more than 1.5 times higher RMRs in the pantropical spotted dolphins would be characteristic of carnivorous marine mammals that adapt to warm-water environments, as observed in the Hawaiian monk seal.

The Kleiber ratios of the Risso’s dolphin in our study (1.5–2.1) was comparable to those observed in beluga whales (1.4–2.2) which are cold-water adapted species in arctic and sub-arctic regions (John et al., 2024). The measurements in beluga whales were conducted using open-flow respirometry with larger individuals (*M*_b_ range: 693–817 kg) than those in Risso’s dolphins in our study (*M*_b_: 315-386 kg). According to the results from our LMM, larger animals have lower Kleiber ratios because of less surface area exposed to water relative to its volume, and the lower minimum value in beluga whales is consistent with this theory, whereas the higher maximum value in beluga whales was not explained by this study. Although the allometric results showed the higher Kleiber ratio in smaller dolphins, measured Kleiber ratio in Risso’s dolphin (1.5-1.9) was higher than those in pantropical spotted dolphins (1.3-1.6). The small number of available individuals at the aquarium was an inevitable limitation in our study, but the LMM analysis with nested random effects of species names mitigated effects of small sample size in each species and showed the metabolic traits as a whole family of Delphinidae with their wide range of body mass (*M*_b_ range: 70-5,300 kg). A potential explanation for the higher Kleiber ratio in Risso’s dolphins as a species level is difference in their cruising speed, in other words, optimal speed. A theoretical study, which divided energy requirements for cost of transport into two factors: metabolic and kinematic costs, predicted higher optimal speed in individuals with higher basal metabolic rate as shown in the following equation: *v* = (BMR/2*k*)^0.33^, where *v* is optimal speed and *k* is a constant (Alexander, 1999; Williams, 1999). Risso’s dolphins are known to have a unique diving behavior of rotating their bodies during descent phases which contributed to an increase of approximately 25% faster in the swim speed than that during non-rotating dives (Visser et al., 2021), but their optimal speed is not clear yet. Short-fined pilot whales are also known to have higher swim speed among deep-diving toothed whales (Soto et al., 2008; Aoki et al., 2017), which tissue samples of locomotor muscles also suggested high metabolic demands with high myoglobin concentrations (Velten et al., 2013). Combining RMR measurements under Kleiber’s criteria with comparisons of muscle morphology and cruising swim speeds in free-ranging Delphinidae species could provide valuable insights into interspecies differences, including the discrepancies observed in our study.

This study showed that RMRs measured by breath-by-breath respirometry were not always lower than those measured by other methods because of the differences in the measuring protocols. Allen et al. (2022) showed that RMRs measured by breath-by-breath respirometry in fasted bottlenose dolphins (141-210 kg) were lower than those reported in other studies (John et al., 2024; a range of *M*_b_: 165-180 kg) and close to the BMR of similar-sized terrestrial mammals, with Kleiber’s ratios of 0.7–1.4. The breath-by-breath respirometry on fasted rough-toothed dolphins (83–123 kg), which are warm-water adapted species in tropical and sub-tropical regions, also reported the Kleiber’s ratios of 1.4–2.2 (Fahlman et al., 2023), similar to that in our study.

Approximately two times of the Kleiber ratio obtained from the largest dolphin measured by breath-by-breath respirometry in our study supports that RMRs measured by both open-flow and breath-by-breath respirometry are comparable as shown in the side-by-side comparison of unfasted resting metabolic rates using the same common bottlenose dolphins by previous studies (van der Hoop et al., 2014; Fahlman et al., 2015).

The *V*_Tinsp_ of a spotted dolphin (ID st_mr) and Risso’s dolphins (ID gg_cf and gg_nf) was 49%, 38%, and 35% of total lung capacity (TLC; TLC = 0.135 *M*_b_^0.92^, Kooyman, 1973), respectively. The *V*_Tinsp_ of Risso’s dolphins was slightly lower than values reported for the bottlenose dolphin (∼45%; Fahlman et al., 2015) and harbor porpoise (∼40%; Reed et al., 2000). A previous study (Fahlman 2024) reported that the resting tidal volume in small and medium-sized cetaceans typically constitutes approximately 30–40% of TLC, which supports that our measurements of tidal volumes fall within the expected range under resting condition.

Assuming similar data quality between the measurement protocols, the RMRs measured using open-flow respirometry from the largest species of Delphinidae, killer whales (Kriete, 1994; Worthy et al., 2013), were included in our allometric analysis. Detailed mechanisms for the smaller mass-specific exponents in Delphinidae compared to those in terrestrial mammals were not examined in this study. Sea otters are known to have the highest mass-specific RMRs among marine mammals due to their small body mass with high metabolic rates (Costa and Kooyman, 1984; Wright et al., 2021). Comparing the lower allometric mass exponents found in an earlier study of sea otters and other marine mammals (Maresh, 2014; Williams, 2022) with those in our study, which focused exclusively on Delphinidae species, further supports the trend of lower mass-specific RMRs in marine mammals with larger body sizes. In the bioenergetic model of cetaceans, the heat loss of animals consists of two factors, body surface and ventilatory fluxes, the ratio of which is estimated to be more than 5-fold higher on the body surface (Sumich, 2021). A morphometric study using 3D modeling found consistency in Bergmann’s rule between two species of pilot whales, in which larger and temperate species of long-finned pilot whales have lower SVR with proportionally larger pectoral flippers compared to those in smaller and tropical or subtropical species of short-finned pilot whales (Adamczak et al., 2020). Negative allometry of pectoral flipper size to body size was also reported in short-beaked common dolphins (Murphy and Rogan, 2006) and killer whales (Clark and Odell, 1999). Although the mass-exponent coefficient with surface area among all of our Delphinidae is not available, a well-fitted mass-specific exponent of 0.66 was also reported in spotted dolphins (*Stenella attenuate*, Worthy and Edwards, 1990). Considering the energy requirements to compensate for heat loss for homeostasis, the majority of the heat loss, which is the proportion of 2/3-power of *M*_b_, may have greater effects on the RMRs of larger marine mammals, making the mass-exponent coefficient lower than 0.75.

This study demonstrated that the scaling effect of body size on RMRs tends to differ from that of terrestrial mammals among Delphinidae with body mass from 70 kg to 5,300 kg. Following the theory of heat loss in water, larger dolphins have lower mass-specific RMRs than that of smaller dolphins do. The magnitude of elevation in RMRs among our warm-water-adapted dolphins was lower than that of other marine mammals, which indicated less metabolic demand for thermoregulation in warmer water than that in colder water. This study filled the knowledge gap of metabolic traits in small- and large-sized dolphins in Delphinidae, but the gap in the smallest size of Delphinidae remained. Future studies conducting metabolic measurements on the smallest Delphinidae, such as Hector’s dolphins (*M*_b_: 40 kg), will contribute to further understanding for both effects of body mass and water temperature on RMRs.

## Ethics statement

All respiratory data were obtained from unrestrained dolphins using routine husbandry training. All procedures, animal husbandry and management were performed under the careful veterinary supervision at the aquarium. The experimental procedures were approved by the Animal Ethics Committee of the Atmosphere and Ocean Research Institute, University of Tokyo (P19-1) and complied with the recommendations of the Life Science Research Ethics and Safety of University of Tokyo for Experiments on Animals.

## Author contributions

Conceptualization: IS, KA, Data-collection: IS, YN, DI, YW, KA, Metabolic rate calculation: KA, Statistical analysis: IS, KA, Writing–original draft: IS, KA, Writing–review and editing: YN, DI, YW, KS, Funding acquisition: IS, YN, KA.

## Acknowledgements

We thank all staff members of the Taiji Whale Museum and Aquarium for their invaluable assistance in handling animals and the measurements of metabolic rate. M. LV and K. Sakai helped with the data collection. A. Fahalman, K. Q. Sakamoto and N. Funasaka provided helpful suggestions. We would like to thank Editage (www.editage.jp) for English language editing.

