## Supplementary material for "Allometric studies on human- and pig-sized Delphinidae to understand resting metabolic rates with respiratory measurements": Figure S1

|  | Measured resting metabolic rate (mL O2 / kg / min) |
| --- | --- |
| Open-flow | 6.5 |
| Breath-by-breath | 6.089 |

**Figure S1.** A side-by-side comparison of unfasted resting metabolic rates in the same bottlenose dolphins (n = 3; identification names of the animals: Kolohe (6JK5), Liko (99L7) and Nainoa (9ON6); a range of body mass: 161-196 kg) by open-flow and breath-by-breath respirometry. Reported values are cited from literature (open-flow respirometry: van der Hoop et al., 2014; breath-by-breath respirometry: Fahlman et al., 2015). There was no significant difference in the measured metabolic rates between the two methods (*p* > 0.05, paired *t*-test). The measurements were conducted in November 2012 for open-flow respirometry and in April 2013 for breath-by-breath respirometry. Over this period, the average water temperature did not differ (25.4˚C in November 2012 and 24.3˚C in April 2013). Bars represent mean ± SD.
